# Patterning effects of FGF17 and cAMP on generation of dopaminergic progenitors for cell replacement therapy in Parkinson’s disease

**DOI:** 10.1101/2024.05.08.593131

**Authors:** Amalie Holm Nygaard, Alrik L. Schörling, Zehra Abay-Nørgaard, Erno Hänninen, Yuan Li, Adrian Santoja, Gaurav Singh Rathore, Alison Salvador, Charlotte Rusimbi, Yu Zhang, Agnete Kirkeby

## Abstract

Cell replacement therapies using human pluripotent stem cell-derived ventral midbrain (VM) dopaminergic (DA) progenitors are currently in clinical trials for treatment of Parkinson’s disease (PD). Recapitulating developmental patterning cues, such as fibroblast growth factor 8 (FGF8), secreted at the midbrain-hindbrain boundary (MHB), is critical for the *in vitro* production of authentic VM DA progenitors. Here, we explore the application of alternative MHB-secreted FGF-family members, FGF17 and FGF18, for VM DA progenitor patterning. We show that while FGF17 and FGF18 both recapitulate VM DA progenitor patterning events, FGF17 induced expression of key VM DA progenitor markers at higher levels than FGF8 and transplanted FGF17-patterned progenitors fully reversed motor deficits in a rat PD model. Early activation of the cAMP pathway mimicked FGF17-induced patterning, although strong cAMP activation came at the expense of EN1 expression. In summary, we identified FGF17 as a promising candidate for more robust VM DA progenitor patterning, with the potential to improve cell products for treatment of PD.

**Summary statement:** In this article we find that FGF17 induces high expression of key dopamine progenitor markers and that these cells can provide functional rescue in a rat model of Parkinson’s disease.

## Introduction

Parkinson’s disease (PD) is an incurable neurodegenerative disorder, the motor symptoms of which are caused by a relatively selective loss of dopamine (DA) neurons in the substantia nigra. Engraftment of VM DA progenitors to the putamen to replace the lost endogenous cells is a promising restorative therapeutic strategy to ensure continuous DA release at physiological levels (Barker and Björklund, 2023), and several approaches are currently in clinical trial (Christiansen and Kirkeby, 2024). The differentiation protocols used to generate VM DA progenitors from human pluripotent stem cells (hPSCs) have been improved by adjusting early patterning with growth factors (Doi et al., 2020; Kim et al., 2021; Nishimura et al., 2023; Nolbrant et al., 2017). Although the current clinically applied differentiation protocols efficiently yield high-purity VM DA progenitor cells, a small proportion of non-VM DA progenitors will inevitably be produced. Fine-tuning of the current protocols to further increase purity and reproducibility is crucial to ensure the success of serial large-scale GMP manufacturing for clinical use.

The isthmic organizer is a key signalling centre located at the midbrain-hindbrain boundary (MHB) of the developing vertebrate embryo (Joyner et al., 2000; Nakamura et al., 2005). It arises from a complex interplay of molecular signals and interactions between neighbouring cell populations and plays a fundamental role in establishing positional information and regional identity of the midbrain and hindbrain. Expression of WNT1 and members of the fibroblast growth factor 8 (FGF8) family are crucial for the function of this secondary organizer in guiding the patterning of the VM in model organisms (Wurst and Bally-Cuif, 2001). Based on this, FGF8 is applied in several protocols for manufacturing of hPSC-derived VM DA progenitors, where it functions to induce expression of the VM DA progenitor marker Engrailed 1 (EN1) (Doi et al., 2020; Nishimura et al., 2023; Nolbrant et al., 2017). High levels of EN1 expression is important for the *in vitro*-generation of *bona fide* caudal VM DA progenitors and correlates with a good graft outcome in animal models of PD (Xu et al., 2000). While *in vitro* protocols have focused only on the use of FGF8, older studies in chick and mouse have shown that not only FGF8, but also FGF17 and FGF18 are expressed at the MHB (Liu et al., 2003). FGF8, FGF17 and FGF18 belong to the same canonical subfamily (FGF8 subfamily), acting on the same receptors (Ornitz and Itoh, 2015), and ectopic application of these FGFs has been shown to stimulate expansion of the midbrain in the developing mouse (Liu et al., 2003). Furthermore, FGF17 has been found to be not only more highly expressed in the developing midbrain region, but also for longer periods of time when compared to both FGF8 and FGF18 (Xu et al., 2000). In humans, however, the dynamics of MHB FGF expression have not been thoroughly explored.

Here, by using single-cell RNA sequencing (scRNAseq) data from an *in vitro* model of the human neural tube (Rifes et al., 2020) and from the developing human fetus (Braun et al., 2023), we identified a broader and stronger expression of FGF17 compared to FGF8 in the human MHB. We further show that FGF17-mediated patterning yielded VM DA progenitors with significantly higher expression of *FOXA2* and *LMX1A* when compared to FGF8. Transplantation of FGF17-derived VM DA progenitors rescued motor deficits in rats and produced DA-rich and highly innervating grafts. Through scRNAseq, we uncovered biological pathways, such as cyclic adenosine monophosphate (cAMP) signalling, underlying the LMX1A upregulation in FGF17-treated cells. In summary, we have identified a new candidate for VM patterning that favours production of the correct population of cells needed for an optimal cell replacement therapy product.

## Results

### Additional FGF8 subfamily members induce VM patterning

Investigation into the FGF expression patterns at the MHB of the developing mouse brain (E11.5), shows that *fgf17* is more broadly expressed around the MHB than *fgf8*, while *fgf18* is only weakly expressed (**Figure 1A)**. A similar broad expression pattern of *FGF17* at the MHB junction was observed in a recently published spatial transcriptomic data of human postconceptional week 5 (pcw 5) fetal tissue (Braun et al., 2023) (**Figure 1B**). In line with this, we have previously found *FGF17* to be expressed at the MHB of an *in vitro* model of the developing human neural tube, and here *FGF17* showed both broader and stronger expression than *FGF8* (**Figure 1C**) (Rifes et al., 2020). While data is not available for *FGF18* in the human embryo at pcw 5, we found *FGF18* to be very weakly expressed in our *in vitro* neural tube model at day 14, aligning with a weak signal in the mouse at E11.5 (**Figure 1A-C**).

**Figure 1.**
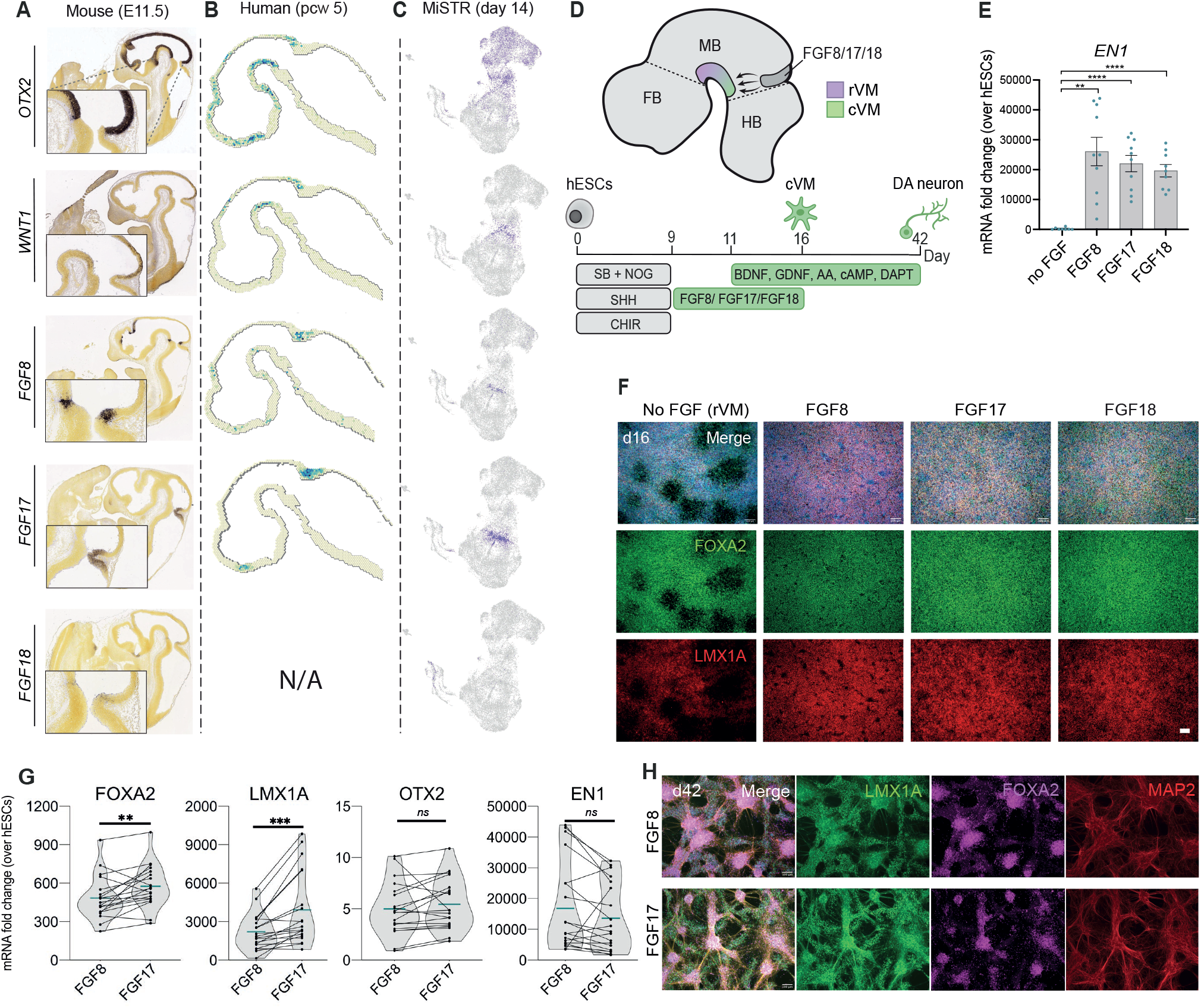
FGF subfamily members are capable of ventral midbrain induction. **A**. Allen Developing Mouse Brain Atlas expression of important MHB genes Otx2, Wnt1, Fgf8, Fgf17, Fgf18 in a E11.5 mouse embryo. **B**. Spatial transcriptomic data of OTX2, WNT1, FGF8, FGF17 in developing human postconceptional week (pcw) 5.5 fetus. **C**. ScRNAseq data from an in vitro human neural tube model (MiSTR) at day 14. **D**. Schematic of expression of endogeneous FGF8 subfamily members at the MHB that induce rostral (rVM) and caudal (cVM) VM patterning as well as schematic of their use in the in vitro protocol. **E**. mRNA expression of EN1 in day 16 VM progenitors treated with No FGF, FGF8, FGF17 or FGF18 analysed by Brown-Forsythe ANOVA followed by Dunnet’s multiple comparison. *p<0.05, **p<0.01, ***p<0.001, ****p<0.0001, ns: non-significant, n=8 (no FGF), n=10 (FGF8, FGF17), n=9 (FGF18). **F**. Immunolabelling of FOXA2, LMX1A in day 16 in vitro derived VM DA progenitors, patterned with either FGF8, FGF17 or FGF18. **G**. Analysis of difference in mRNA expression of important VM DA progenitor markers between FGF8 and FGF17. **p<0.01, ***p<0.001, EN1 p= 0.11, ns: non-significant, analysed with a paired t-test, n = 19. **H**. Immunolabelling of FOXA2, LMX1A and MAP2 in day 42 in vitro derived DA neurons. All scalebars: 100µM.

To compare the ability of different MHB-associated FGFs from the FGF8 subfamily to fine-tune hPSC derived VM DA progenitors, we added either FGF8, FGF17 or FGF18 from day 9 to 16 of an already published directed differentiation protocol for the production of clinical-grade VM DA progenitors (Kirkeby et al., 2023; Nolbrant et al., 2017), and differentiated the cells in parallel (**Figure 1D**). By qRT-PCR analysis we found that all three FGFs induced a significant increase in the caudal VM DA marker *EN1 mRNA* expression levels (**Figure 1E**) while maintaining cultures with a high purity of FOXA2+/LMX1A+ progenitors (**Figure 1F**). Through qRT-PCR analysis of parallel batches of VM differentiations, we found that FGF17 induced significantly higher expression of *FOXA2* and *LMX1A* compared to FGF8, while *OTX2* and *EN1* expression did not differ (**Figure 1G**). We found no difference in the expression of these markers between FGF17- and FGF18-patterned batches (**Figure S1A**). Given the higher in vivo relevance of *FGF17* over *FGF18* at the MHB (**Figure 1A-C**) we therefore decided to focus on the actions of FGF17 for the remainder of the study. Nuclear flow cytometry of FGF17-patterned day 16 VM DA progenitor cells showed unchanged proportions of FOXA2/OTX2 double-positive cells compared to paired FGF8-patterned cells (**Figure S1B**), indicating that the positive effect of FGF17 on FOXA2 expression may in part be due to increased expression levels within each cell.

### FGF17-patterned VM DA progenitors reverse motor deficits in a rat model of PD

To assess the maturation capacity of FGF17-treated VM DA progenitors, we performed *in vitro*-maturation, and found that both FGF17- and FGF8-patterned VM DA progenitors produced cultures of neurons expressing TH, MAP2, FOXA2 and LMX1A at day 42 of differentiation (**Figure 1H and S1C**). To ascertain whether the FGF17-patterned VM DA progenitors were also functional upon engraftment, FGF17-patterned VM DA progenitors were transplanted to the striatum of unilaterally, 6-hydroxydopamine (6-OHDA)-lesioned nude rats (**Figure 2A**). Immuno-histochemical assessment showed dense human neural fiber-innervation of the rat striatum (**Figure 2B**) and DA neuron-rich grafts (**Figure 2C**). The transplanted cells yielded full amelioration of motor deficits at 27-weeks post-transplantation in the amphetamine-induced rotation test (**Fig. 2D**). We next confirmed that the FGF17 grafts expressed markers of the A9-subtype of DA neurons through co-labelling of TH with the floorplate marker FOXA2 (Kee et al., 2017; Kirkeby et al., 2017) and the A9 DA neuron marker ALDH1A1 (Carmichael et al., 2021) (**Figure 2E**). The FGF17-patterned grafts generated on average 2335±812 mature TH^+^ neurons per 1e5 transplanted cells (mean±SD). This is comparable to the yield obtained from the clinical-grade FGF8-patterned VM DA progenitor cell product STEM-PD, which is currently being tested in a European phase-I/IIa clinical trial for PD, ClinicalTrials.gov number NCT05635409 (2835±1466 TH^+^ cells, **Figure 2F**, data from Kirkeby et al., 2023).

**Figure 2.**
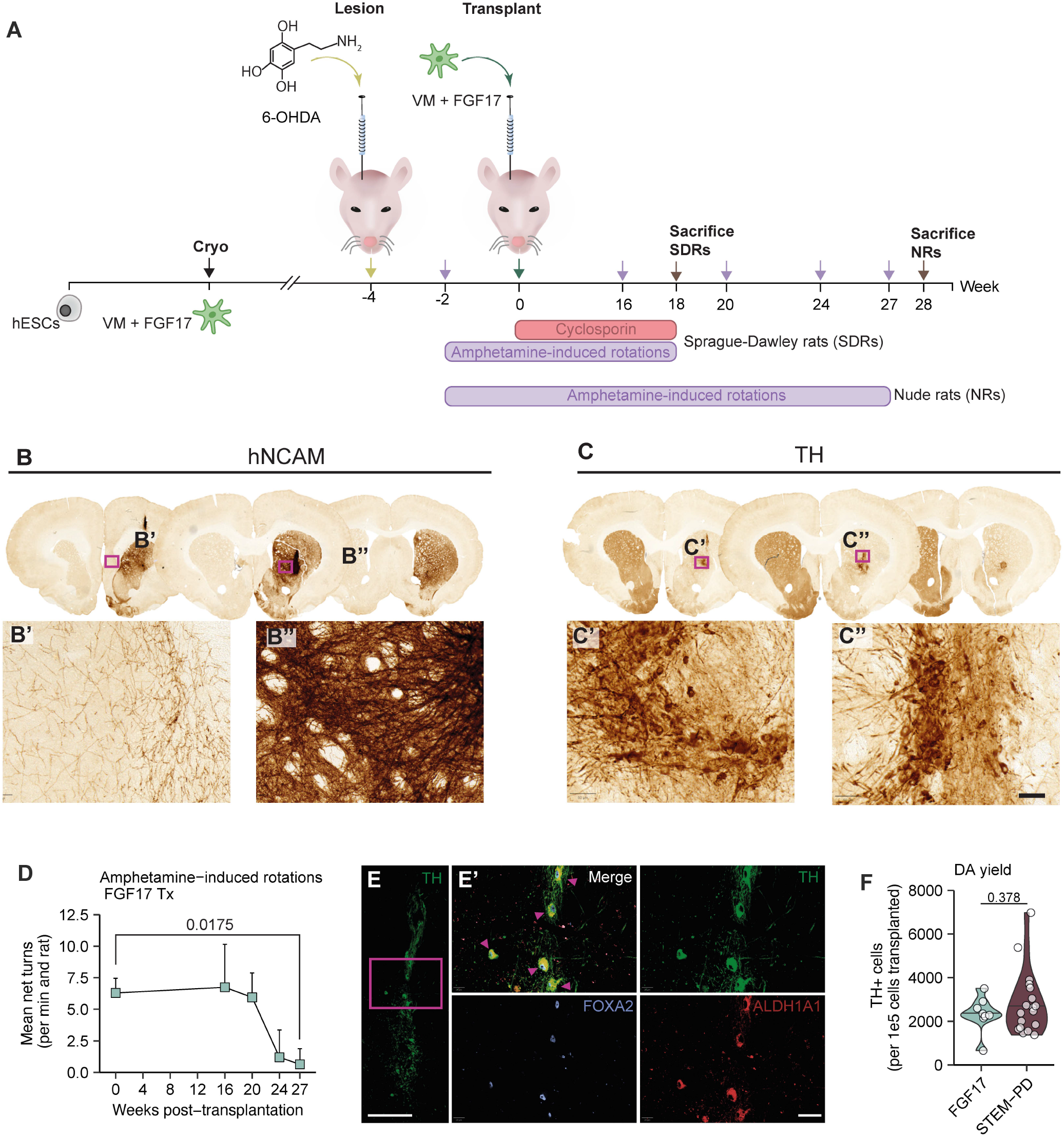
Functionality of FGF17-patterned DA neurons upon transplantation to Parkinsonian rats. **A**. Experimental outline: Nude rats (NRs) and Sprague-Dawley rats (SDRs) were unilaterally lesioned with 6-OHDA and then transplanted with cryopreserved day 16 FGF17-patterned cells. The functionality of the transplanted cells was evaluated at 16-, 20-, 24- and 27-weeks post-transplantation through amphetamine-induced rotations. SDRs were immunosuppressed with Cyclosporin upon transplantation. **B-C**. Representative images of immunohistochemical stainings for hNCAM (B) and TH (C) in transplanted rat brains, showing dense human fiber innervation and TH+-rich grafts. **D**. Mean net turns per min in amphetamine-induced rotation test from rats transplanted with FGF17-patterned cells. n=7 (week 0), n=4 (week 16, 20, 24), n=3 (week 27). A Welch t-test was used to determine differences between week 0 and 27 (p-value = 0.0175). **E**. Representative images of FGF17 graft at 27 weeks, immunofluorescently labelled for TH, FOXA2 and ALDH1A1. Triplepostive cells are indicated by triangles. Overview scale bar = 200 µm, zoom-in scalebar = 40 µm. **F**. Quantified total yield of TH+ cells per 1e5 cells transplanted in FGF17 grafts at 27 weeks compared to the clinical cell product STEM-PD. Data for STEM-PD obtained from Kirkeby et al., 2023. n=8 (FGF17), n=18 (STEM-PD). Statistical significance determined by t-test, p=0.378.

### RNA-sequencing reveals temporal differences in FGF8 and FGF17 signalling

To uncover the biological pathways underlying the differences observed between FGF8- and FGF17-patterned cells, we performed two different RNA sequencing studies: 1) A time-course bulk RNA-seq experiment on day 9 VM progenitors treated with either FGF8 or FGF17 for up to 24 hours; and, 2) A scRNAseq experiment on hashtag-labelled day 16 VM DA progenitors patterned with either FGF8 or FGF17 for 7 days, from day 9-16 (**Figure 3A)**. For the bulk RNA-seq, differences in FGF response pathways were investigated at the earliest time points immediately after initiation of FGF treatment (i. e. for 15 min, 60 min, 4 hours or 24 hours after adding the FGFs, n=3 for each FGF). Principal component analysis showed that at 15 min post-FGF addition, there were negligible differences between the groups, but that the groups differed transcriptionally at the 1, 4 and 24 hour time-points (**Figure 3B**). Differentially expressed gene (DEG) analysis at 1-4 hours post FGF treatment showed a strong induction of a whole group of ERK/MEK-pathway associated early response genes (*EGR2, EGR3, EGR4, IER2, IER5, ARC* and *FOSB*). Interestingly, all of these genes were more upregulated in FGF8-treated cells compared to FGF17-treated cells (**Figure 3C-D)**.

**Figure 3.**
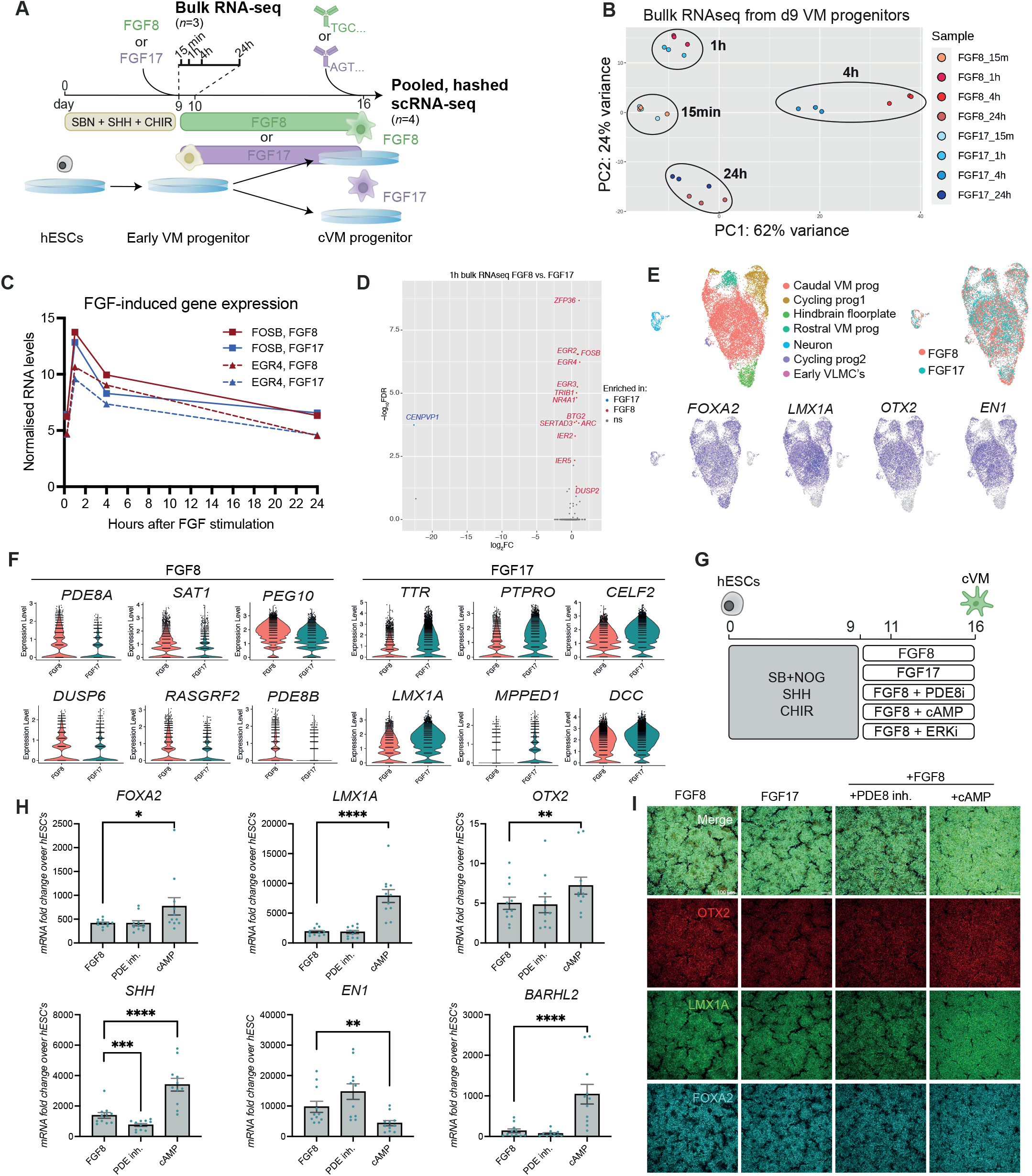
Addition of cAMP during VM patterning mimicks the effect of FGF17. **A**. Experimental outline for bulk and scRNAseq. FGF8- or FGF17-patterned day 9 VM DA progenitors were (1) collected at 15 min, 1h, 4h, or 24h after FGF addition for bulk RNAseq (n=3 per condition), or (2) differentiated until day 16 caudal ventral midbrain progenitors for scRNAseq, n=4. **B**. PCA plot comparing bulk RNAseq of FGF8 and FGF17 treated VM DA progenitors at all four timepoints, n=3. **C**. Expression trajectories of early response genes after addition of FGFs (here represented by FOSB and EGR4). **D**. Volcano plot of DEGs at 1hr between FGF8- and FGF17-treated VM DA progenitors. **E**. UMAP projection of scRNAseq data from FGF8 and FGF17 day 16 VM DA progenitors. **F**. Violin plots of top DEGs between FGF8 and FGF17 cells in the scRNAseq data. **G**. Outline of in vitro differentiation strategy to test target pathways uncovered by RNAseq. **H**. Quantitative RT-PCR of key VM markers in day 16 progenitors treated with either PDE8i or cAMP together with FGF8 from day 9-16. One-way ANOVA for paired data followed by Dunnet’s multiple comparisons test was performed to analyse differences in gene expression *p<0.05, **p<0.01, ***p<0.001, ****p<0.0001, ns: non-significant, n=11. **I**. Immunocytochemistry showing co-staining of OTX2, LMX1A and FOXA2 on day 16 VM DA progenitors treated wih PDE8i or cAMP, scalebar: 100µM.

Next, we performed scRNAseq on hashtag-labelled day 16 VM DA progenitors patterned with either FGF8 or FGF17 (n = 4 differentiation experiments, each experiment testing FGF8 and FGF17 in parallel, **Figure 3A**). All 8 batches of differentiated cells showed similar transcriptional profiles at day 16 (**Figure S2**), affirming the high similarity between VM batches differentiated with either FGF8 or FGF17. As expected, the majority of cells sequenced produced a large cluster of caudal VM DA progenitors marked by expression of *FOXA2, LMX1A, OTX2* and *EN1* (**Figure 3E**). Both treatments also gave rise to a small proportion of rostral VM progenitors, marked by the absence of *EN1* and presence of *OTX2, LMX1A* and *FOXA2*. Furthermore, we found a small cluster of contaminating hindbrain floorplate progenitors (*FOXA2*^+^ but *OTX2*^-^) in both treatment groups, arising primarily from one of the 4 differentiation experiments, VM#4 (**Figure 3E, S2B**). Analysis of the top DEGs between all FGF8- and FGF17-treated cells revealed enrichment of *PDE8A, PDE8B* and *DUSP6* in FGF8-treated cells. The PDE8 family members are phosphodiesterases that play a major role in the hydrolysis of cAMP and *DUSP6* (MKP3) is a protein phosphatase that has previously been found to be coexpressed with FGF8 at the MHB (Echevarria et al., 2005). FGF17-treated cells in contrast expressed higher levels of the thyroid binding hormone, transthyretin (TTR) and protein tyrosine phosphatase receptor type O (PTPRO) within the FOXA2+/LMX1A+/OTX2+/EN1+ VM DA progenitor cluster (**Figure 3F and S2)**. PTPRO has recently been shown to be enriched on DA progenitor cells (Xu et al., 2022).

To assess the impact of the targets identified in the DEG analysis on progenitor patterning, we tested addition of commercially available agonists and antagonists of the pathways identified in the bulk and scRNAseq analyses in combination with FGF8 treatment (**Figure 3G**). Based on the difference in ERK-related signalling between FGF8 and FGF17 treated cells seen in the bulk RNAseq experiment, we tested an ERK inhibitor (ERKi) from day 9 to 16 together with FGF8 treatment from day 9 to 16. While we observed no effect on the expression of *FOXA2* and *LMX1A*, we found a significant decrease in *EN1* expression when ERKi was maintained from day 9 to 16 (**Figure S3A**). In addition, we found expression of the diencephalic marker *BARHL2* to be significantly increased when cells were treated with ERKi. The phenotype observed in the ERKi-treated cultures resembled that of cells in the ‘No FGF’ group. These results show that ERK is necessary for FGF8 mediated induction of *EN1* expression as well as repression of the diencephalic marker *BARHL2*.

We next proceeded to test addition of membrane-permeable cAMP (dibutyryl-cAMP, referred to as cAMP) as well as a PDE8 inhibitor (PF-04957325, PDE8i) to the cultures, as the scRNA-seq data indicated involvement of the cAMP pathway in the different action between FGF17 and FGF8. Addition of cAMP is normally used in the late-stage of DA protocols (after day 16) to induce neuronal maturation (Doi et al., 2014; Kim et al., 2021; Nishimura et al., 2023; Nolbrant et al., 2017), but in these experiments we tested cAMP as well as the PDE8i together with FGF8 earlier in the protocol, from day 9-16. Surprisingly, early addition of cAMP to FGF8-treated cultures caused a significant increase in expression of VM markers *FOXA2, LMX1A, OTX2* and *SHH* by day 16 of differentiation (**Figures 3H-I and S3B**), thereby exacerbating the phenotype observed for FGF17-compared to FGF8-patterned progenitors (**Figure 1F**). This indicated that increased activation of the cAMP pathway may be a main mechanism differing between FGF8 and FGF17 signalling. However, early treatment with cAMP also induced a significant decrease in expression of the caudal VM marker *EN1* as well as an upregulation of the diencephalic marker *BARHL2*, implying that the cells generated with early addition of cAMP were of a more rostral VM identity which is not beneficial for DA generation (**Figure 3H and S3B)**. Interestingly, treatment with a PDE8i, which would be expected to increase endogenous levels of cAMP, did not yield the same effects as cAMP in the VM DA progenitors (**Figure 3H-I and S3B)**. This implies that the VM DA progenitors are not at this stage producing high levels of cAMP endogenously.

Finally, to uncover more details on the signalling dynamics of FGF8 versus FGF17, we employed the computational tool Domino (Cherry et al., 2021) to identify intercellular signalling and transcription factor activation, specifically in our ‘caudal VM prog’ cluster (**Figure S3C)**. Among the most important networks expressed in the FGF8-treated caudal VM progenitor cluster were components of the TGF and PDGF pathway, governed by *EN1* and *FOXA1*. In the FGF17-treated cluster, signalling through the neurotrophin ligand receptor *NTRK3* was predicted to be related to *LMX1A* expression. To investigate a potential role of these pathways we tested PDGFR inhibition or activation and the addition of neurotrophin 3 (NT-3) in combination with FGF8 treatment. However, by day 16 of differentiation we observed no significant difference in mRNA levels of key VM DA markers between the treatment groups, and concluded that these pathways were not crucial for caudal VM DA patterning (**Figure S3D-F**).

We have previously shown that the addition of FGF8 to the cell differentiation medium fine-tunes cell fates to ensure a higher yield of VM DA progenitors compared to the rostrally adjacent subthalamic nucleus progenitors, which do not yield DA neurons after terminal maturation (Kirkeby et al., 2017). However, the use of other FGF8 subfamily members remains largely neglected, dispite their recognition at the MHB decades ago. At the turn of the century, a collection of studies on MHB development demonstrated that FGF submembers FGF8 and FGF17 could act synergistically during early MHB patterning (Picker et al., 1999; Reifers et al., 1998, 2000; Xu et al., 2000). Overall, our data indicates that both FGF17 and FGF18 can work independently of FGF8 to govern VM DA patterning, and that the effect of these three FGFs are to a large degree redundant. This is in line with the knowledge that FGF8 subfamily ligands all function on the same receptors (Olsen et al., 2006). Not surprisingly, we could confirm that the effect of FGF8 is highly dependent on the activity of ERK.

Despite the similar patterning effects, minor differences in activated signalling pathways between FGF8 and FGF17 were evident. Although the protocols for patterning of VM DA progenitors are already very efficient, even minor improvements to purity can be crucial for successful product release when manufacturing cells in a GMP setting for clinical use. The transcriptional differences between FGF8 and FGF17 resulted in the identification of the cAMP pathway as a pathway which, when activated early, can increase expression of VM genes LMX1A and FOXA2. However, this comes at the expense of the crucial VM DA progenitor marker EN1, which we have shown to be required for producing successful DA grafts (Kirkeby et al., 2017). Interestingly, another VM DA progenitor product which is currently in clinical trial for PD by BlueRock Therapeutics includes cAMP in the early patterning of their VM DA progenitors, from day 10-16 (Kim et al., 2021). Based on this, it would be relevant to investigate if there is a minimal threshold of EN1 expression required at the progenitor stage for achievement of optimal in vivo efficacy.

In summary, we have herein demonstrated the ability of several FGF family members to induce VM patterning, and dissected the potential mechanisms by which specifically FGF17 might enhance early MHB cell fate decisions. The use of FGF17 might be benefical compared to FGF8 for producing an efficacious better stem cell product for transplantation, ultimately additionally ameliorating PD patients’ motor symptoms. The observation that FGF8 +PDE8 inhibition has the opposite effect of FGF8+cAMP on expression levels of key VM DA progenitor markers opens for future investigations into the underlying mechanism of this discrepancy and whether transplantation of PDE8 inhibitor or cAMP-treated VM DA progenitors can achieve functional recovery in an animal model of PD.

## Author contributions

A.K. conceived the study. A.H.N., A.S., Ad.S. and C.R. performed *in vitro* differentiations. Z.A. and A.H.N. performed cell hashing and ICC. E.H., G.S.R and Y.L. performed bioinformatic analyses. A.S., Y.Z. and A.L.S. performed in vivo transplantations. A.L.S. performed behavioral tests and graft characterization. Z.A. compiled data and A.K., Z.A., A.H.N. and A.L.S. wrote the manuscript.

## Acknowledgements

We thank the 10X sequencing platform SCOP at the University of Copenhagen as well as the Microscopy and Genomics platforms at reNEW Copenhagen.

## Data availability

Spatial data from human 5 pcw fetal tissue is available at https://github.com/linnarsson-lab/developing-human-brain/ and can be accessed through https://doi.org/10.1126/science.adf1226. Developing mouse brain data is available from Allen Developing Mouse Brain Atlas, https://developingmouse.brain-map.org. Data from the in vitro model of the developing human neural tube (MiSTR) can be accessed through https://www.ncbi.nlm.nih.gov/geo/query/acc.cgi?acc=GSE135399. scRNAseq and bulk RNAseq datasets generated specifically for this study will be accessible upon publication.

## Materials and methods

### Cell culturing

Mycoplasma-free, karyotyped RC17 (Roslin cells) hESCs were cultured on Lam521 (1mg/ml) coated plates. RC17 were cultured in StemMACS iPS Brew XF (Miltenyi Biotec) and passaging was performed using 0.5mM EDTA dissociation at 37°C. For the first 24h after passaging 10µM ROCK inhibitor (Miltenyi Biotec) was included in the media. Specification of hESC towards VM fates was performed in accordance with the Nolbrant et al., 2017 protocol. Adaptation to the current protocol was the addition of either FGF8b (100 ng/ml, #423-F8-025/CF), FGF18 (100ng/ml, #8988-F18-050) or FGF17b (100ng/ml, #319-FG-025) to test for optimal VM DA patterning. FGFs were in all cases sourced from R&D Systems to avoid any comparibility issues related to the vendor. Isoform b was chosen for FGF17 due to its potency and analogy to FGF8b. FGF18 does not undergo alternative splicing. Cells were harvested at day 16 and day 42 for analysis.

For the additional factor screening experiments VM patterned cells were treated with PDE8 inhibitor PF-04957325 300nM (MedChem, #HY-15426), dibutyryl-cAMP 500uM (Sigma-Aldrich, #D0627), DUSP6 inhibitor (E/Z)-BCI hydrochloride 100ng/ml (Sigma-Aldrich, #B4313), T3 40ng/ml (Sigma-Aldrich, #T6397), PDGFR inhibitor CP673451 100nM (R&D Systems, #CP 673451 5993/10), PDGFC 1ug/ml (R&D Systems, #1687-CC-025) or NT-3 10-20ng/ml (R&D Systems, #267-N3-005) on top of FGF8b 100ng/ml from day 9 to 16. Analysis was performed at day 16 and 42. In experiments with ERK inhibition cells were treated with Trametinib 50nM (Biotechne #7709/10) day 9 to 16 on top of FGF8b 100ng/ml from day9 to 16. Analysis was performed on day 16.

### Animal experiments

All procedures were conducted in accordance with the European Union Directive (2010/63/EU) and was approved by the local ethical committee at Lund University and the Swedish Department for Agriculture (ethical permit number 5.2.18-10992/18). Adult, female Sprague Dawley (SD) and athymic, nude rats (Hsd:RH-*Foxn1*^rnu^) were purchased from Charles River and Envigo, respectively, and were housed on a 12:12-hr light:dark cycle with *ad libitum* access to food and water, n = 4 SD rats, n = 7 nude rats. For all surgical procedures, rats weighting >225 g and/or older than 3 months were anesthetized via intraperitoneal (IP) injection of a 20:1 or 3:2 mixture of fentanyl-Dormitor or ketaminol-Dormitor (Apoteksbolaget), respectively, according to the weight. The DA neurons of the rats were unilaterally ablated by intracranial injection of 10.5 µg 6-hydroxydopamine (6-OHDA) to the medial forebrain bundle (MFB). The extent of the lesion was assessed by amphetamine-induced rotations test. For this purpose, 3.5 mg/kg amphetamine was administered by IP injection, and mean net turns per min were measured over 90 min, following a 10 min staggered start. Cryopreserved VM DA progenitors derived from RC17 hESCs were prepared as previously described (Kirkeby et al., 2012; Nolbrant et al., 2017). 3e5 cells were unilaterally transplanted to SD- and nude rat striatum as described previously (Tiklová et al., 2020). SD rats were injected at the coordinates AP, +0.8; ML, -2.8; DV, -4.5, and nude rats at the coordinates AP, +0.9/+1.4; ML, - 3.0/-2.6; DV, -5.0/-4.0. SD rats were administered 10 mg/kg cyclosporine by daily IP injections two days prior to transplantation, and until euthanization, 18 weeks post-transplantation.

### Immunolabelling

Brains were fixed in 4% (wt/vol) paraformaldehyde by perfusion, according to standard protocol. The brains were removed, and post-fixed over-night before dehydration in 25% (wt/vol) sucrose. Brains were sectioned with a thickness of 35 µm in 1:8 series by using a freezing microtome (Leica), and were stored in **Buffer A** (13 mM NaH_2_PO_4_, 38 mM Na_2_HPO_4_ 30% (vol/vol) ethylene glycol, 30% (vol/vol) glycerol) at -20°C. Immunolabelling was performed on free-floating sections, using the Corning® Netwells® system (Merck). The sections were placed on a slowly rotating (∼100 rpm) orbital shaker during all incubations, which were done at room temperature, unless specified otherwise. Sections were washed for 3x 5 min in PBS between all steps unless otherwise specified. For antigen retrieval, sections were incubated in **Buffer B** (10 mM Tris Base, 1 mM EDTA Solution, 0.05% Tween 20, pH 9.0, H_2_O) at 80°C for 30 min. During immunohistochemistry – where the detection system was horseradish peroxidase (HRP)-based – endogenous peroxidases were inactivated by incubation in **Buffer C** (10% (vol/vol) methanol, 3% (vol/vol) H_2_O_2_, PBS) for 30 min. If necessary to decrease background toning in IHC, sections were first blocked in **Buffer D** (2.5% vol/vol Triton-X, 5% (vol/vol) species-specific serum), supplemented avidin and biotin, using the Avidin/Biotin Blocking Kit (Vector Laboratories) according to the manufacturer’s instructions. Sections were blocked in **Buffer D** for 1h. The sections were after this incubated with one or several of the primary antibodies ALDH1A1 (1:1000, AbCam, ab52492), hNCAM (1:1000, Santa Cruz Biotechnology, Sc-106), TH (1:2000, Merck Millipore, AB152), TH (1:1000, Merck Millipore, AB1542), HuNu (1:1000, Merck Millipore, MAB1281), FOXA2 (1:1000, Santa Cruz Biotechnology, sc-101060) and LMX1A (1:1000, Merck Millipore, AB10533) over-night. The sections were then blocked for 15 min in **Buffer D**. After this, the section were incubated in the biotinylated secondary antibody or fluorescently labelled secondary antibodies for 1h for IHC and immunofluorescence (IF), respectively. For IHC, the sections were thereafter incubated in ABC HRP or ABC alkaline phosphatase complex according to the manufacturer’s instructions (Vector Laboratories). The chromogenic substrate development in IHC was thereafter performed by incubating the sections in the enzymatic system-compatible substrate solution until the desired colour intensity had developed. The following substrate were used: (i) 0.5 mg/mL DAB, catalysed by the addition of 0.125% (vol/vol) H_2_O_2_, (ii) Vector® Blue or (iii) Vector® DAB, with or without supplementation with nickel (Vector Laboratories), prepared according to the instructions. Sections were then immediately washed several times briefly in PBS to avoid colour saturation. IF necessary during IHC, counterstaining was done using a mild progressive labelling with haematoxylin. Chromagenically-labelled sections were mounted and dehydrated by incubation in step-wise-increasing concentrations of ethanol and were then cleared. Lastly, the sections were coverslipped in an appropriate mounting media. Cells were fluorescently labelled with the primary antibodies (**Table S2**) as previously described (Nolbrant et al., 2017).

### Imaging and quantification

The rat brains were processed and sectioned in series of 8, and one series was used for the quantifications. For quantification of TH^+^ cells in DAB-labelled grafts, brightfield, Z-stack images under 20X magnification were captured and stitched automatically by using a Leica DMI6000B with the LAS-X software program. TH^+^ cells in FGF17 grafts were counted manually in the software program Fiji (version 2.1.0) whereas the TH^+^ cells in the STEM-PD graft were counted manually at the microscope. The quantifications of TH^+^ cells is presented as estimated total number of TH^+^ cells per 1e5 cells transplanted per animals using the formula:

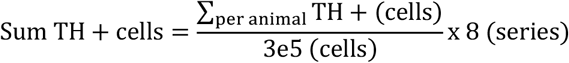

Overview, 10X brightfield images of the chromagenically labelled rat brains was acquired on a Axio Scan Z1 (Zeiss) miscroscope. IF images were acquired using the 40X objective on the confocal microscope Stellaris (Leica). Brightfield and IF images were processed in the software program QuPath (version 0.4.3). Imaging of immunolabelled cells was performed on the Leica AF600 widefield epifluorescence microscope, using Leica LAS-X software. Images were processed with Fiji (version 2.1.0).

### Quantitative RT-PCR

RNA was collected on day 16 and 42 from 500,000 cells. For RNA isolation the RNeasy Micro Plus kit (Qiagen) was used. cDNA synthesis was performed from 1ug purified RNA using the Maxima First strand synthesis kit (ThermoFisher, #K1642). SYBR green (Roche, #04887352001), primer mix (fwd+rev) and cDNA for each qRT-PCR reaction were pipetted using the liquid dispensing robot iDOT (Dispendix). Ct values were measured over 40 cycles on the Lightcycler 480 (Roche). Each reaction was run in duplicates and the average Ct value were used for calculating fold change values over hESCs by the 2^-ΔΔCt^ method (Livak and Schmittgen, 2001).

### Single cell RNA sequencing

Four batches of paired FGF8- and FGF17-treated day 16 VM DA progenitors (8 samples total) were thawed in wash media containing DMEM/F12 + 5%KOSR. Next, cells were counted, spun down and resuspended in PBS(wo Ca and Mg) with 0.5%BSA at 5mio cells per ml. 0.5mio cells per sample were stained with 0.5µg of unique cell surface hashing antibody (Biolegend) and incubated for 30 min at 4 degrees. After antibody tagging, cells were washed three times in PBS(wo Ca and Mg) with 0.5%BSA and resuspended at 1000 cells/ul in PBS(wo Ca and Mg) +0.5%BSA. The eight samples were prepared to be run across two 10X sequencing lanes by pooling two batches of paired FGF8 and FGF17 samples at 1:1:1:1 ratio for both sequencing lanes. Each sequencing lane were loaded with 25,000 cells. For sequencing of our day 16 VM DA progenitors, we combined 10X scRNA-seq with CITE-Seq for sample multiplexing to sequence eight different replicates across two protocols (Stoeckius et al., 2018). We performed cell surface hashing on 4 different batches per treatment group and each 10X lane ran 2x FGF8 and 2 xFGF17 treated groups to correct for batch effect. The raw sequence data has been preprocessed with the Alevin-fry pipeline (He et al., 2022) and the downstream analyses were performed with Seurat v.4.1.0. The acquired count matrices were filtered to remove low quality cells and doublets. All mitochondrial and ribosomal genes were removed from the dataset before continuing with normalization by SCTransform. The dimensionality of the data was reduced with PCA and UMAP, followed by graph-based clustering and cluster-wise differentially expressed gene identification. Differentially expressed gene analysis between the FGF8 and FGF17 protocols were performed using DESeq2 v.1.30.1(Love, 2021) as implemented in the FindAllMarkers function of Seurat v.4.10. Genes with *p_val_adj* < 0.05 were accepted as significantly differentially expressed genes. In order to compare the signalling network between the two studied protocols, we analyzed the datasets for the two protocols separately with identical parameters. We first identified regulons (modules of genes that are targeted by transcription factors) and quantified regulon activity in each cell using Pyscenic v.0.11.2 (Van de Sande et al., 2020). We then used Domino v.0.1.1 (Cherry et al., 2021) to infer the signalling networks. Transcription factors that were specific to each cluster were identified using the Mann-Whitney U test, top transcription factors (up to ten) with p-values below 0.001 were selected. We next identified potential receptors that activated transcription factors, where receptors and transcription factors were considered significantly connected if Pearson correlation>0.23 with a maximum of ten receptors for each transcription factor. Afterwards, we acquired the ligand signalling partners for each connected receptor from the CellphoneDB2 database. Ligands that were not expressed were excluded. With the top ten transcription factors identified for each cluster and their connected receptors and ligands, we generated the cluster-specific top regulon activity/transcription factor activation heatmap, the transcription factor and receptor correlation heatmap, the global signalling network, as well as the global intercellular signalling network.

### Bulk RNAseq

RC17 cells were patterned towards VM according to Nolbrant et al. 2017. On day 9, cells were treated with either FGF8b 100 ng/mL or FGF17 100 ng/mL for 15 min, 1 h, 4 h and 24 h. Upon sample collection cells were washed once with cold PBS -/- and lysed in RLT buffer (QIAGEN). RNA was extracted using RNeasy Mini kit (QIAGEN) including on column DNase treatment (QIAGEN). RNA quality was assessed using the HS RNA screentape kit (Agilent) on Tapestation 4200 (Agilent) following manufacturers protocol. Samples with RIN^e^ values >8 were included for next steps. Libraries were prepared using the NEBNext Poly(A) mRNA magnetic isolation module (NEB #E7490S) in combination with NEBNext Ultra II directional RNA library prep kit for Illumina (NEB #E7765) following manufacturer’s protocol. Libraries were eluted in 17.5ul 0.1X TE after the final clean up. Library yields were quantified using the Qubit and quality control were performed on Tapestation 4200 (Agilent) using HS D1000 screentape kit (Agilent) following manufacturer’s protocol. Libraries were diluted to 10 nM and pooled. One additional clean up were performed on the pooled sample to eliminate an adaptor contamination in one sample. All 24 libraries were sequenced together on one flow cell on Illumina NextSeq 2000. The obtained sequencing data underwent quality control and alignment with the nf-core/rnaseq (v3.10.1) (Patel et al., 2024) pipeline. Default parameters were used, except for setting skip_trimming to true, and hg38 was used as reference genome for the alignment. To ensure the data and alignment quality, we inspected the MultiQC report from the pipeline. The uncorrected gene count matrix generated by the pipeline was quality-filtered by removing genes with low counts across samples. The resulting count matrix served as an input for differential expression analysis, performed using the DESeq2 package (v1.40.2) (Love et al., 2014), to identify genes with differential expression patterns between FGF8 and FGF17 treatments at different time points of stimulation. To visualize the differences between the treatments the results from differential expression analysis and the normalized gene counts were visualized using a volcano plot and a heatmap.

### Spatial gene expression visualisation

Publicly available spatial transcriptomics data from post-conception week 5 human neural tube (Braun et al., 2023) was used to visualize spatial gene expression in developing human brain. The spatial data was converted into an AnnData object, a format required by the Tangram (Biancalani et al., 2021) package. To create the figures, we modified the plotting functions from Tangram to suit our purposes.

### Statistical analysis

Statistical tests and visualization were done by using R (version 4.2.2) or Prism (10.0.02), with a critical significance level of alpha ≤ 0.05. Equality of variance was investigated with Brown-Forsythe tests, and normality with Shapiro-Wilk tests and manual inspection of frequency distributions. Homoskedastic and normal data was analysed with the omnibus test analysis of variance (ANOVA), and data not meeting these criterions with a Kruskal-Wallis test. Post-hoc analysis was done by using a Dunnet’s multiple-comparison test. Comparison of two groups with normal distributions was done with paired or non-paired two-sided t-tests, with consideration for equality of variance. Unless otherwise specified, the data is expressed as mean ± SEM.

