## Supplementary material for "Patterning effects of FGF17 and cAMP on generation of dopaminergic progenitors for cell replacement therapy in Parkinson’s disease"

### Supplementary tables

**Table S1.** List of primers.

| Gene | Full name | Primer sequence (fw/rv) |
|---|---|---|
| **ACTB** | Beta-actin | CCTTGCACATGCCGGAG/ GCACAGAGCCTCGCCTT |
| **BARHL1** | BarH-like homeobox 1 | GTACCAGAACCGCAGGACTAAA/ AGAAATAAGGCGACGGGAACAT |
| **BARHL2** | BarH like homeobox 2 | GGAGATTACGAGTAGCCGTGAG/ AAGCTACGCTCCAGTTGATTGA |
| **CORIN** | Corin, serine peptidase | CATATCTCCATCGCCTCAGTTG/ GGCAGGAGTCCATGACTGT |
| **EN1** | Engrailed 1 | CGTGGCTTACTCCCCATTTA/ TCTCGCTGTCTCTCCCTCTC |
| **FOXA2** | Forkhead box A2 | CCGTTCTCCATCAACAACCT/ GGGGTAGTGCATCACCTGTT |
| **FOXG1** | Forkhead box G1 (BF1) | TGGCCCATGTCGCCCTTCCT/ GCCGACGTGGTGCCGTTGTA |
| **GAPDH** | Glyceraldehyde-3-phosphate dehydrogenase | TTGAGGTCAATGAAGGGGTC/ GAAGGTGAAGGTCGGAGTCA |
| **HOXA2** | Homeobox A2 | CGTCGCTCGCTGAGTGCCTG/ TGTCGAGTGTGAAAGCGTCGAGG |
| **LMX1A** | LIM homeobox transcription factor a | CGCATCGTTTCTTCTCCTCT/ CAGACAGACTTGGGGCTCAC |
| **NKX2-1** | NK2 homeobox 1 | AGGGCGGGGCACAGATTGGA/ GCTGGCAGAGTGTGCCCAGA |
| **OTX2** | Orthodenticle homeobox 1 | ACAAGTGGCCAATTCACTCC/ GAGGTGGACAAGGGATCTGA |
| **PAX6** | Paired box 6 | TGGTATTCTCTCCCCCTCCT/ TAAGGATGTTGAACGGGCAG |
| **PITX2** | Paired like homeodomain 2 | AACTCTATGAACGTCAACCCCC/ CGACATGCTCATGGACGAGATA |
| **SHH** | Sonic hedgehog | CCAATTACAACCCCGACATC/ AGTTTCACTCCTGGCCACTG |
| **TH** | Tyrosine hydroxylase | CGGGCTTCTCGGACCAGGTGTA/ CTCCTCGGCGGTGTACTCCACA |
| **WNT1** | Wingless-type MMTV integration site family, member 1 | GAGCCACGAGTTTGGATGTT/ TGCAGGGAGAAAGGAGAGAA |

**Table S2.** List of reagents and the dilutions.

| Reagent | Dilution | Manufacturer | Cat. # |
|---|---|---|---|
| ALDH1A1 | 1:1000 | AbCam | ab52492 |
| hNCAM | 1:1000 | Santa Cruz Biotechnology | Sc-106 |
| TH | 1:2000 | Merck Millipore | AB152 |
| TH | 1:1000 | Merck Millipore | AB1542 |
| HuNu | 1:1000 | Merck Millipore | MAB1281 |
| FOXA2 | 1:400 | R&D Systems | AF2400-SP |
| FOXA2 | 1:1000 | Santa Cruz Biotechnology | sc-101060 |
| LMX1A | 1:1000 | Merck Millipore | AB10533 |
| MAP2 | 1:1000 | Sigma | M1046 |
| OTX2 | 1:500 | R&D Systems | AF1979 |
| EN1 | 1:1000 | Novo Nordisk A/S | N/A |
| Anti-Goat Cy3 | 1:200 | Jackson ImmunoResearch | 705-165-147 |
| Anti-mouse 488 | 1:200 | Jackson ImmunoResearch | 715-545-150 |
| Anti-Rabbit 647 | 1:200 | Jackson ImmunoResearch | 711-605-152 |
| Horse anti-mouse | 1:200 | Vector Laboratories | BA-2001 |
| Rabbit anti-sheep | 1:200 | Vector Laboratories | BA-6000 |
| Goat anti-rabbit | 1:200 | Vector Laboratories | BP-9100 |
| Haematoxylin Gill No. II | - | Sigma Aldrich | GHS232-1L |

### Supplementary figures

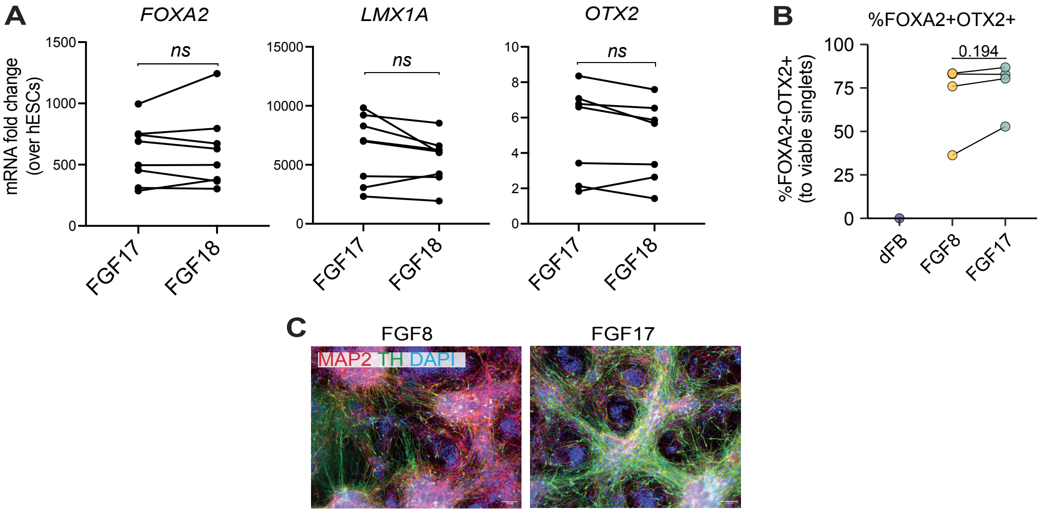


**Figure S1. Comparison of FGF8- and FGF17-patterned VM DA progenitors**
**A.** Comparison of mRNA expression of key VM markers in FGF17- and FGF18-treated day 16 progenitors analysed by a paired t-test. ns: non-significant, n=7. **B.** Flow cytometric analysis of FOXA2/OTX2 double-positive cells. Paired t-test, p= 0.194, n=4. **C.** Immuolabelling of day 42 FGF8- and FGF17-treated cells.

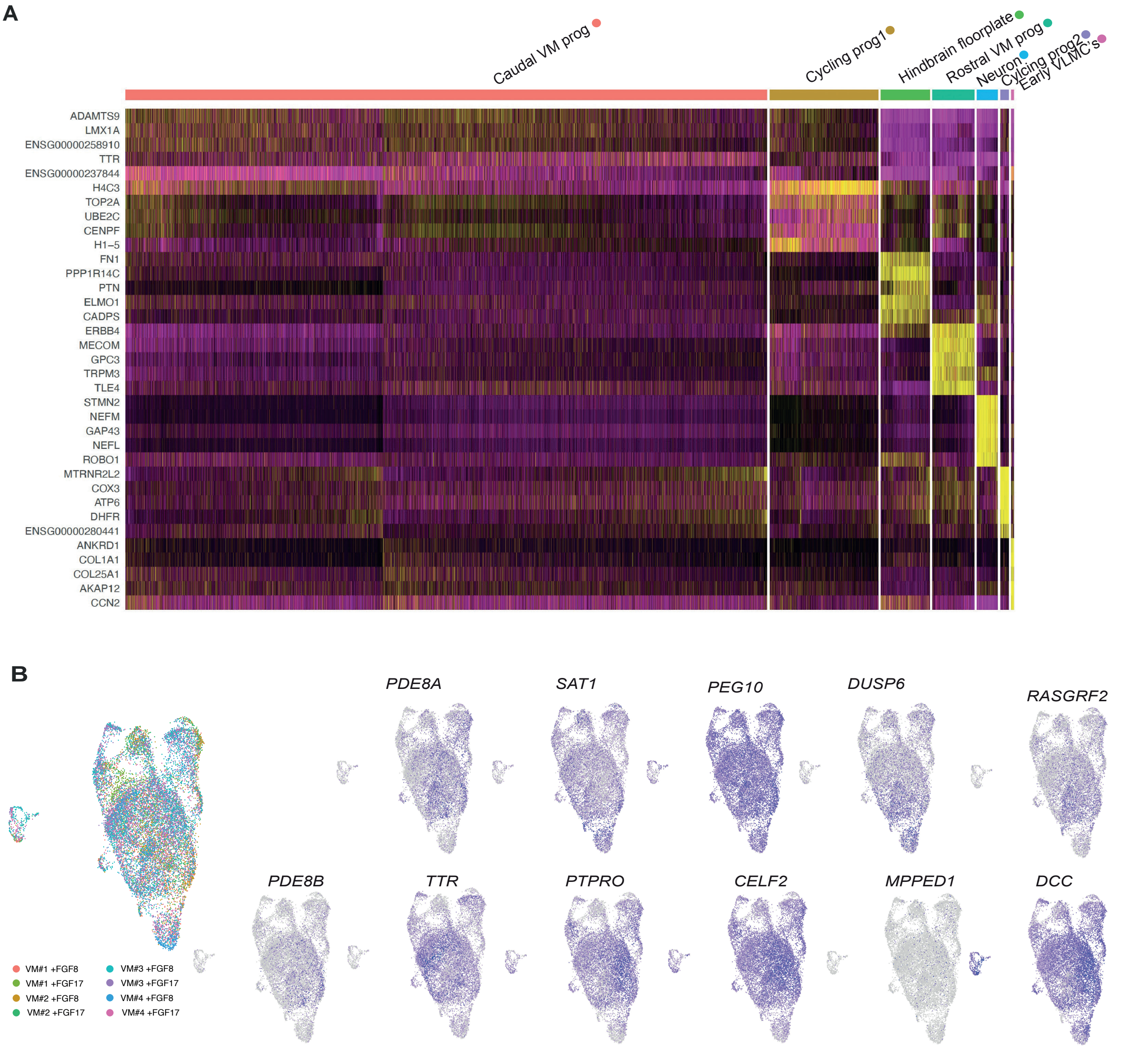


**Figure S2. Gene expression in day 16 FGF8- and FGF17-patterned progenitors**
**A.** Heatmap showing DEGs in each of the annotated clusters in the day 16 FGF8+FGF17 scRNAseq dataset. **B.** Feature plots in UMAP space for genes found to be differentially expressed between FGF8- and FGF17-patterned progenitors on day 16 (from Fig. 3E). Top left plot shows hashtags specific to each biological replicate included in the experiment.

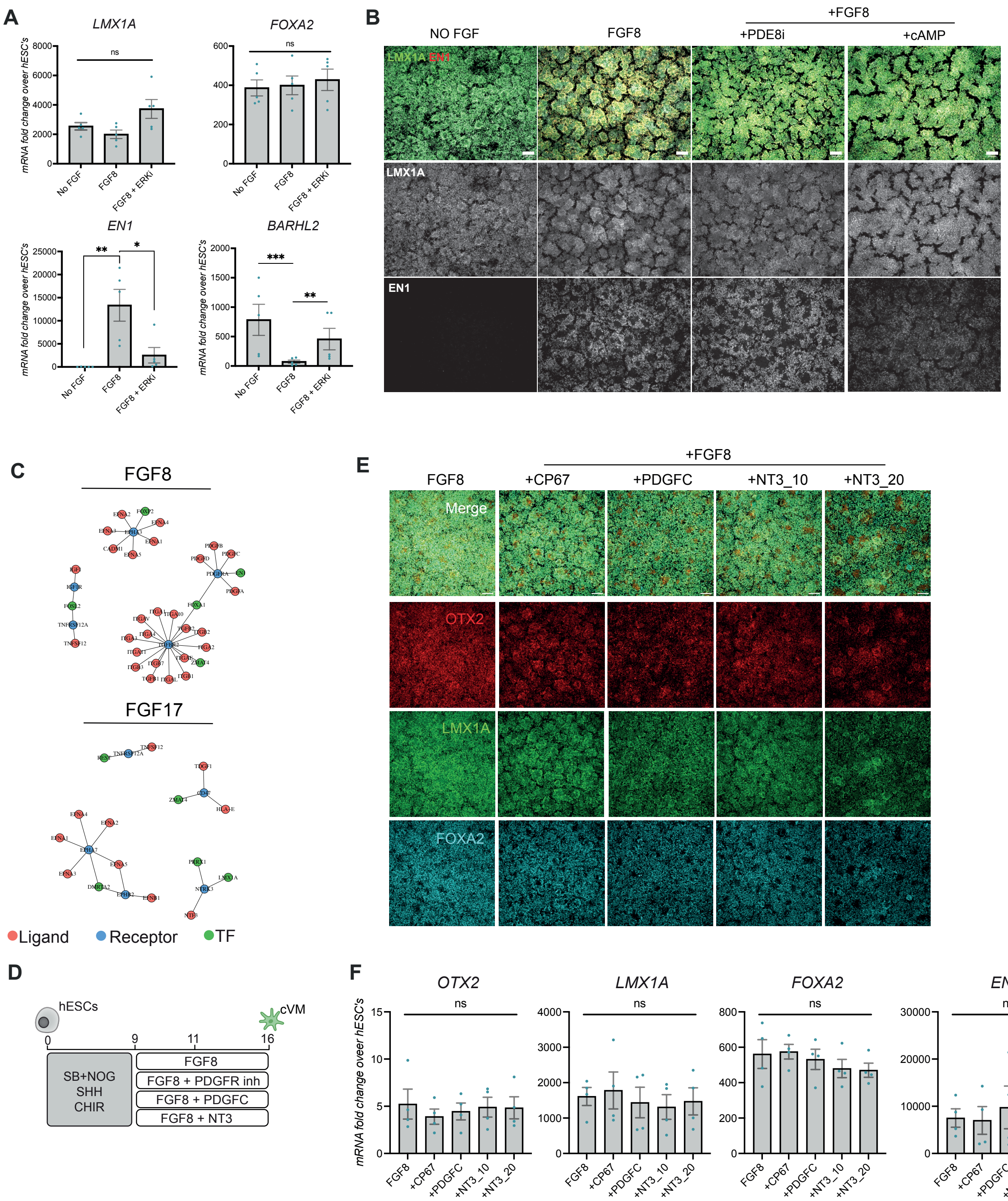


**Figure S3. Investigation of signalling pathways in FGF8- and FGF17-mediated patterning**

**A.** Quantitative RT-PCR of VM markers in day 16 progenitors treated with an ERK inhibitor (Trametinib) from day 9-16 in combination with FGF8. Difference in expression between groups were tested using One-way ANOVA for paired data followed by Dunnet's multiple comparisons test. $^*p<0.05$, $^{**}p<0.01$, $^{***}p<0.001$, ns: non-significant, n=5. **B.** ICC of day 16 VM DA cultures treated with either PDE8i (PF-04957325) or cAMP together with FGF8 from day 9-16, scalebar = 100μM. **C.** Intercellular signalling network of the "caudal VM progenitor" cluster from either FGF8-treated or FGF17-treated cultures in the day 16 scRNAseq data, using Domino. **D.** Schematic of testing conditions performed on the basis of Domino results. **E.** ICC of day 16 VM DA cultures treated with either PDGFR inhibitor (CP67), PDGFC or NT-3 at 10 or 20 ng/ml, all in combination with FGF8 from day 9-16, scalebar = 100μM. **F.** Quantitative RT-PCR of key VM markers in day 16 progenitors treated with the same compounds as in E. One-way ANOVA for paired data followed by Dunnet's multiple comparisons test or Friedman test followed by Dunn's multiple comparison were used to test differences in gene expression, ns: non-significant, n=4.
